# A geometrical framework for *f* –statistics

**DOI:** 10.1101/2020.08.19.257709

**Authors:** Gonzalo Oteo–García, José–Angel Oteo

## Abstract

A detailed derivation of the *f*–statistics formalism is made from a geometrical framework. It is shown that the *f*–statistics appear when a genetic distance matrix is constrained to describe a four population phylogenetic tree. The choice of genetic metric is crucial and plays an outstanding role as regards the tree–like–ness criterion. The case of lack of treeness is interpreted in the formalism as presence of population admixture. In this respect, four formulas are given to estimate the admixture proportions. One of them is the so–called *f*_4_–ratio estimate and we show that a second one is related to a known result developed in terms of the fixation index *F*_*ST*_. An illustrative numerical simulation of admixture proportion estimates is included. Relationships of the formalism with coalescence times and pairwise sequence differences are also provided.

## 1 Introduction

In the course of time a population will be subject to drift, migration, natural selection, etc., whose recurrence eventually leads to the branching of subgroups as new populations with distinct allele frequencies. However, this might not always be the case. To characterize the *tree–like–ness* or lack of this, different measures have been introduced, for example *f*–statistics [10, 7, 8, 6, 12].

We build up a geometrical framework where the *f*–statistics emerge as the constraints of a genetic distance matrix to correspond to a phylogenetic tree of four populations. The idea is to consider populations as points in an allele frequency space whose dimension equals the number of SNPs in the dataset and to interpret the three statistics *f*_2_, *f*_3_, and *f*_4_ of the *f*–formalism as scalar products of differences of allele frequency vectors. This approach allows also to point out some limitations of the *f*–formalism.

An *f*–statistics based criterion for tree–like–ness or lack of it is discussed and interpreted in geometric terms in the allele frequency space. In the population admixture case, four formulas are derived to estimate the admixture proportions. One of them is the already known *f*_4_–ratio result and another one proves to be similar to a result in [4] developed in terms of the fixation index *F*_*ST*_ [14]. Eventually, a relationship of *f*–formalism with two results in [8] based on coalescence times and pairwise sequence differences is established.

In Section 2 the geometrical framework is introduced. The algebraic constrains on a distance matrix to be related to a phylogenetic tree are studied in Section 3 and the subsequent derivation of *f*–statistics formalism is in Section 4. The formulation of the tree–like–ness criterion is given in Section 5. Section 6 contains applications of the geometrical approach in a treeness scenario and points out some limitations of the *f*–formalism. The population admixture case is treated in Section 7. Relationships with coalescence times, pairwise differences and fixation index *F*_*ST*_ are given in Section Section 9 contains the discussion and conclusions. Two technical Appendices are included.

## 2 Allele frequency space *S*_*f*_

Every population *P*_*i*_, with *i* = 1, …, *n*, may be characterized by an array that can be represented by a point, or a vector 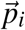 of dimension ℝ^*m*^, which is the number of loci considered. For population *P*_*i*_ we denote

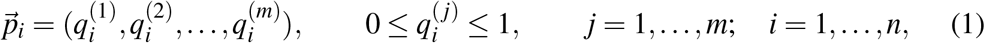

where 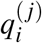 is the allele frequency of SNP *j* in population *P*_*i*_. Every vector is bounded by 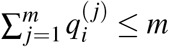.

Populations that are genetically close to each other appear as clustered points in this space. As a consequence of genetic evolution this picture has a dynamic character because the numerical values of frequencies 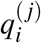 do change in time. The net effect is a variation in the modulus and orientation of vector 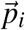 which describes an erratic trajectory in ℝ^*m*^ as time passes. Figure 1a gives a representation of the situation in ℝ^3^. At an initial time three populations are represented by 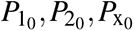. At a later time they are found in *P*_1_, *P*_2_, *P*_x_, due to genetic evolution. The crinkly lines are mere illustrations of the paths they might have followed in the space.

**Figure 1:**
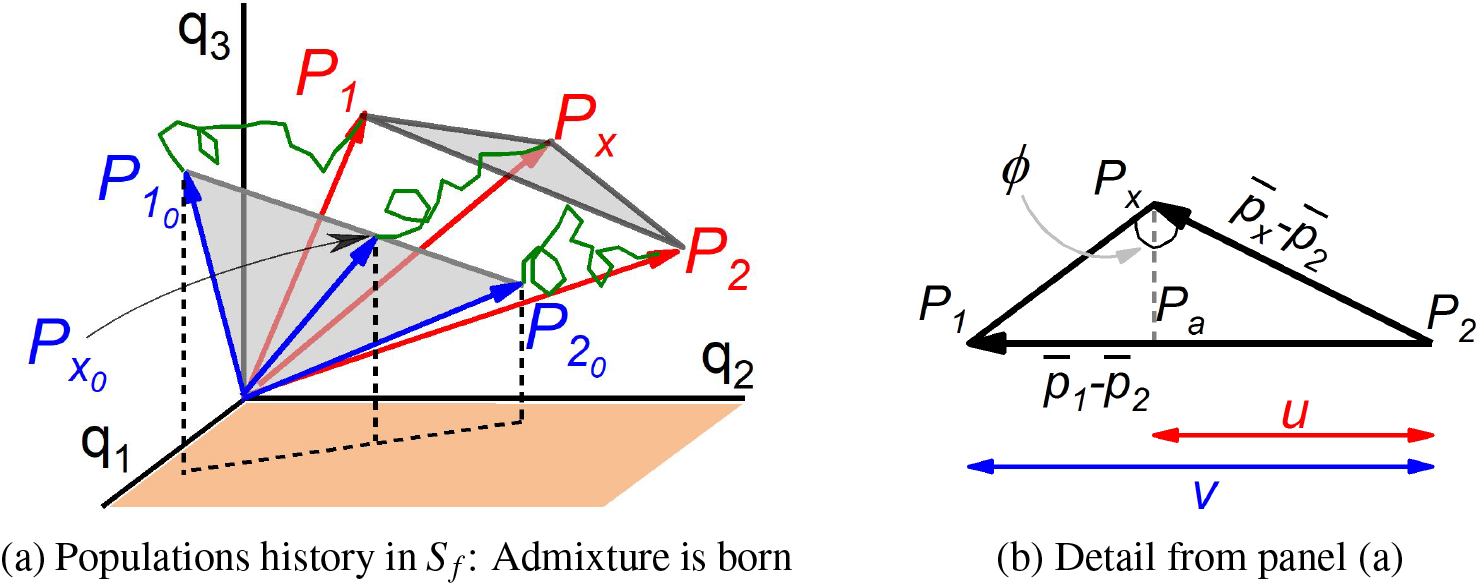
(a) Three–dimensional representation of the birth of the admixed population 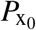 from 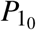 and 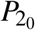, in *S*_*f*_. As time passes the three vectors describe erratic trajectories (green) and give rise to evolved populations *P*_1_, *P*_x_, *P*_2_: The segment 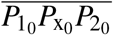 becomes triangle *P*_1_*P*_x_*P*_2_. (b) Enhanced illustration of the triangle in panel (a). The ratio of lengths *u* over *v* gives the admixture proportion estimate *α*_*ls*_ in (30).

Given three population vectors 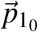, 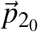, 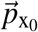 in ℝ^*m*^, we will make extensive use of the following linear combination

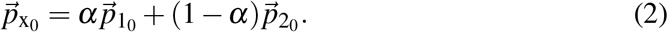

where 0 < *α* < 1. It means that we build up a new population vector 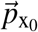 out of 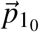 and 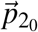, such that every vector component *i* is obtained as: 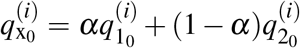. Figure 1a illustrates an important property: The three vectors involved in the linear combination are coplanar. Besides, 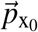 is the inner vector in the triplet and its tip stays on the segment 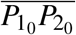. Moreover, the ratio obtained from length of segment 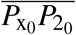 over length of segment 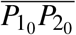, equals the value *α*. These features will be important to asses the nature of the approximations we will build up to determine a population mixing proportion.

One further technical element that we need to introduce is the ordinary Euclidean scalar product (or dot product) of two vectors, defined as

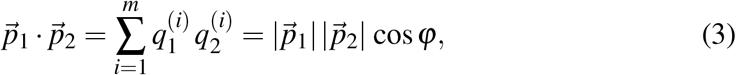

with 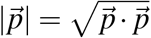, the modulus of vector 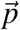; and φ the angle between the two vectors.

Let us assume that vector 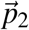 is of unit length, i.e. 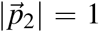. Then 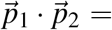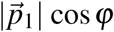, which is the length of the orthogonal projection of vector 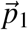 along the direction defined by the unit vector 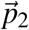. This geometric interpretation of the scalar product and the two (equivalent) definitions of the dot product (3) will be instrumental in the interpretation of *f*–formulas.

We will denote *S*_*f*_ the *allele frequency space* ℝ^*m*^ endowed with the operations linear combination and scalar product.

Eventually, the choice of a particular genetic distance in *S*_*f*_ will be a crucial point in the derivation of the *f*–statistics formalism. The mathematical definition of distance has a number of requirements. We will pay attention to the so–called *triangle inequality* or *sub–additivity property* which establishes that the distance in *S*_*f*_ between *P*_*i*_ and *P*_*j*_ is always smaller or equal to the sum of distances between *P*_*i*_ and *P*_*k*_, on one side, and *P*_*k*_ and *P*_*j*_ on another.

## 3 Genetic distance matrix and treeness

Given a set of *n* populations in *S*_*f*_, it is always possible to express part of the information in terms of a symmetric *genetic distance matrix* 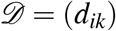, where *d*_*ik*_ stands for a genetic distance measure between populations *P*_*i*_ and *P*_*k*_, with *i, k* = 1, …, *n*. For the time being we work with a generic genetic distance measure.

We commence by working out a formal result in [11] to provide the explicit algebraic conditions needed for 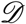 to correspond to a phylogenetic tree, in the cases with three and four populations. Concomitantly, an important idea introduced by *f*–statistics formalism consists in studying trees with three and four populations to avoid direct analysis of the whole putative tree, which is in general a cumbersome task. These results lead in a simple way to introduce the *f*–statistics as algebraic constraints on the distance matrix with a particular metric.

### 3.1 The elements of the four–population tree

The four tips and outer branch lengths in Figure 2a are denoted *P*_*i*_ and *ℓ*_*i*_, (*i* = 1, 2, 3, 4), respectively. The inner branch is denoted *ℓ*_0_. Besides, the distances between populations *P*_*i*_ and *P*_*k*_ are represented by the matrix elements *d*_*ik*_. For the time being no particular metric is chosen. The following system of six linear equations and five unknowns is the algebraic formulation of the tree topology

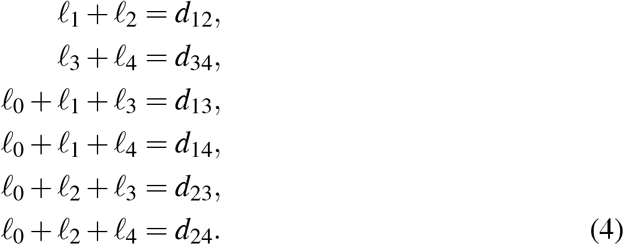

**Figure 2:**
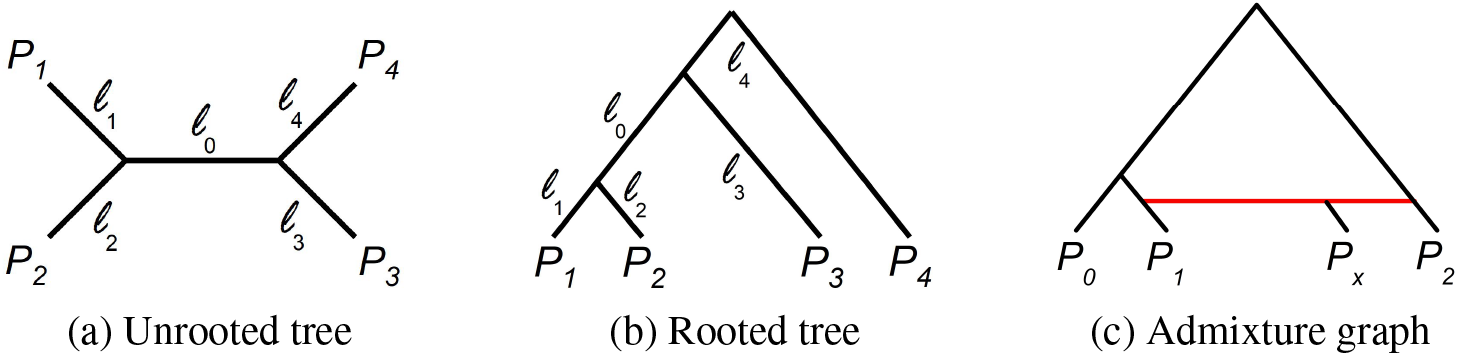
Four–population (*P*_*i*_) trees with branch lengths (*ℓ*_*i*_) in (a) and (b). Admixture graph (c) where populations *P*_1_, *P*_2_ give rise to *P*_x_. The length of admixture branches (red) is proportional to the contributor weight.

According to [11, Lemma 1] a necessary and sufficient condition for the matrix 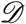 to have a tree realization is that the system above has a solution in non–negative numbers. Higher dimension cases are solvable in terms of this one. The condition for system (4) to have a solution requires the augmented matrix of system (4)

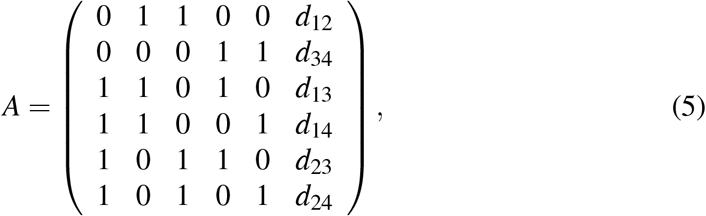

to be rank(*A*) = 5. This condition implies det(*A*) = 0, and after some algebra yields the algebraic treeness condition

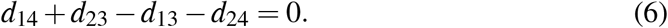

In such a case the solution

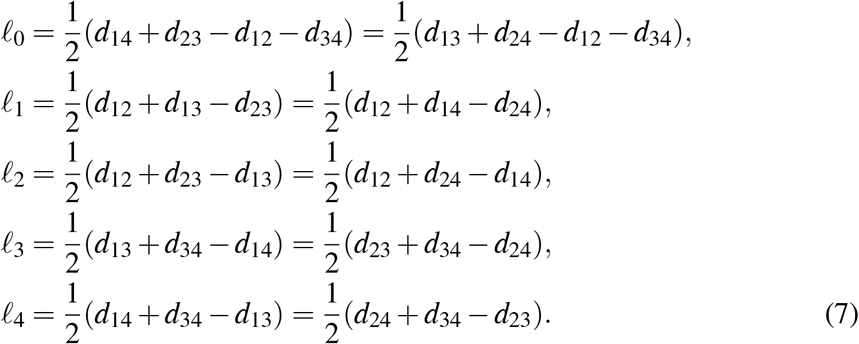

Once evaluated, if all *ℓ*_*i*_ are non–negative numbers then the tree exists and is unique. More specifically, given that all *d*_*ij*_ ≥ 0, then *ℓ*_0_ ≥ 0 in the first line of (7) implies

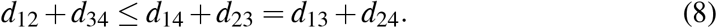

Alternatively, if (8) holds then it is *ℓ*_0_ 0 in (7). This inequality is known as *four–point condition* [1] and guarantees, on its own, the tree to be unique. Observe that it merely establishes that the inner branch length *ℓ*_0_ is a positive quantity.

Due to the symmetry of the distance matrix 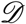 and the equation system, only the trees (((*P*_1_, *P*_2_), *P*_3_), *P*_4_), (((*P*_1_, *P*_3_), *P*_2_), *P*_4_) and (((*P*_1_, *P*_4_), *P*_2_), *P*_3_), have to be considered [11].

So far no mention has been made to the particular distance definition used to build up the matrix 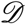 and therefore all these results are generic. Any mathematical distance among the plethora of existing definitions is valid, although some of them might capture more information from the dataset at hand than others.

The equation system (4) already appears in [2, Figure 79] where three different approaches are suggested: Least squares, maximum likelihood and minimum evolution. Interestingly, it is the algebraic solution developed above which leads to the *f*–statistics framework, as we will show.

### 3.2 The elements of the three–population tree

The algebraic solution of a three–population tree goes as follows. There are three unknowns, the branch lengths *ℓ*_1_, *ℓ*_2_, *ℓ*_3_, and three equations for the tree topology

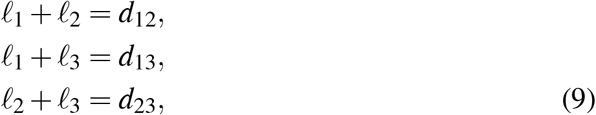

whose solution reads

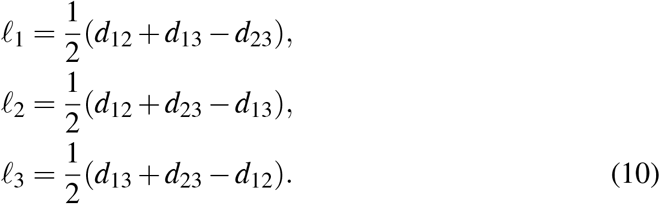

The positive–ness of branches, *ℓ*_1_ > 0, *ℓ*_2_ > 0, *ℓ*_3_ > 0, should be guaranteed by the sub–additivity property of the genetic distance. However, it turns out in some practical cases that the numerical value of one branch is negative. All we can say then is that those three populations are not related by genetic drift, at least as regards the distance definition used. Hence, the lack of treeness is related to the breaking of the triangle property of the distance.

## 4 Derivation of *f*–statistics from a genetic distance matrix

The *f*–statistics formalism is readily obtained from the framework developed above when the following genetic distance is chosen

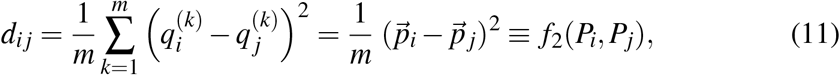

where we have introduced the notation *f*_2_ to refer to the matrix elements of 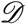 with this choice of metric. The definition (11) corresponds to a squared Euclidean distance, a fact that requires important mathematical considerations. We address this issue bellow in Section 5.

The four population algebraic treeness condition (6) becomes the following *f*–formula

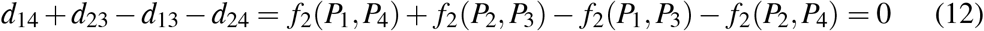

The sum of *f*_2_ terms in this equation can be expressed as a single scalar product which, in turn, leads to the definition of the statistic *f*_4_

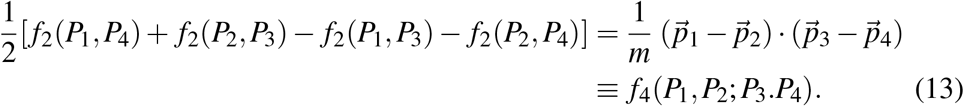

The treeness condition (12) reads then

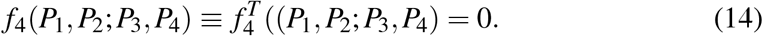

The inner branch length *l*_0_ is provided by (7) and may be written down in terms of *f*_4_ too

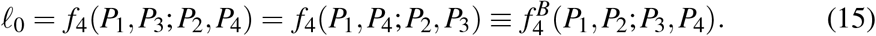

Following [8], we have introduced the superscript notation with 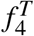, for *treeness* test; and 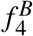, for inner *branch* length evaluation; in order to stave off explicit rearrangements in the arguments of *f*_4_. Thus, *f*_4_ serves two purposes in the context of treeness. The evaluation of 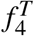 proceeds directly as in definition (13), whereas the computation of 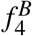 is preceded by an argument reordering.

The lengths of the tree external branches provided by (7) have the structure: *d*_*ij*_ + *d*_*ik*_ − *d*_*jk*_, which should be a positive quantity according to the triangle inequality. Interestingly, it can be shaped as a scalar product

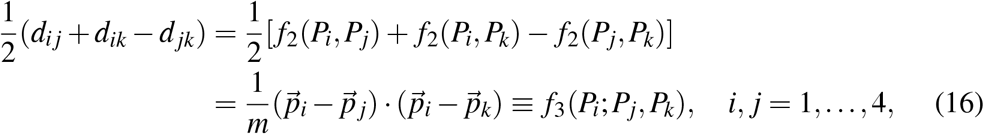

where we have defined the new statistic *f*_3_. Therefore

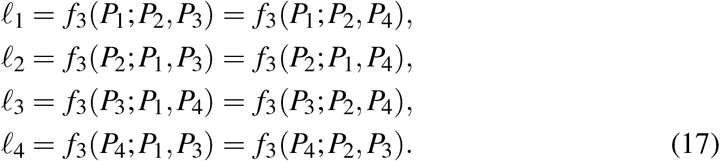

In Appendix B we gather a number of useful algebraic *f*–identities.

Figure 3 illustrates the geometric meaning of treeness condition (12): For the given tree (black) the sum of lengths in red, on one side, and in blue, on another, must match. Alternatively, the four point condition (8) can be also used. In terms of *f*_2_ it reads

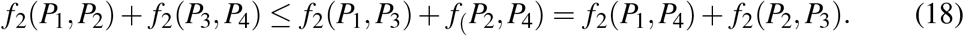

**Figure 3:**
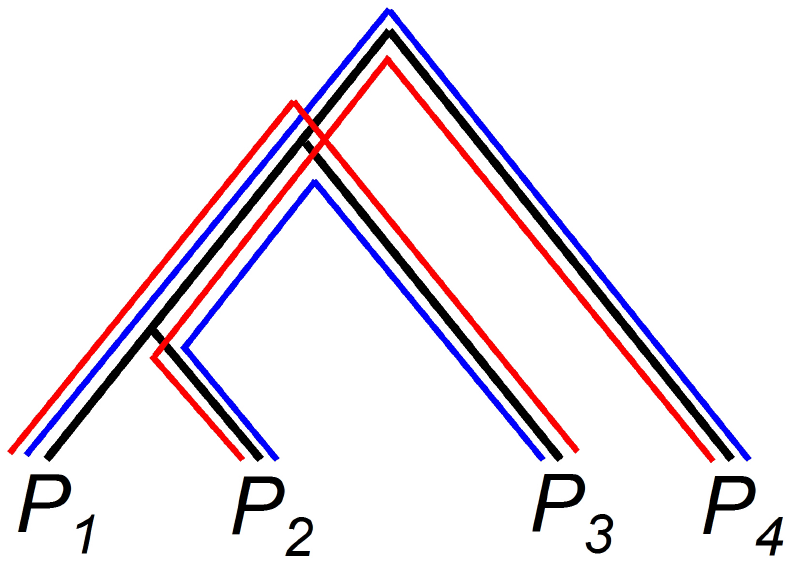
With four populations, treeness condition (12) requires the total length in red to equate that in blue.

Eventually the equations (10) for the three–population tree branch lengths become

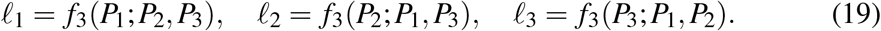

## 5 The metric *f*_2_: Triangle criterion

As defined in (11), *f*_2_ is a squared Euclidean distance in *S*_*f*_, because it is the scalar product of a vector with itself, scaled by the space dimension. However, the squared Euclidean distance is not properly a metric because the triangle inequality (or sub–additivity property) that establishes *d*_*ik*_ ≤ *d*_*ij*_ + *d*_*jk*_, for all *i, j, k*; does not hold for any combination of three vectors. It is however possible to add some constraints that may render *f*_2_ a true distance in some practical situations. To see this let us consider three arbitrary population vectors 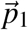, 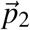, 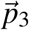, in *S*_*f*_ and work out the scalar product that appears in the definition of 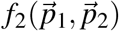

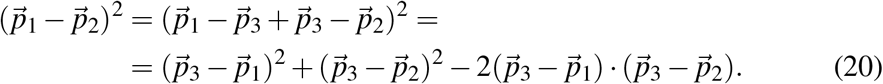

Using (11) and (16) we get the equivalent expression

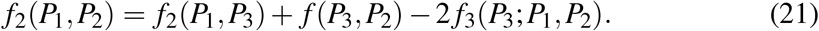

If and only if the scalar product in the rhs of (20) is positive, or *f*_3_(*P*_3_; *P*_1_, *P*_2_) > 0 in (21), the sub–additivity condition is preserved. According to the definition of scalar product, the condition 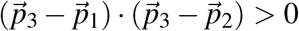, means that the angle *ϕ* between the difference vectors 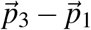 and 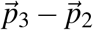, lays in the interval *ϕ* ∈ [0, 90°]. Since sub–additivity must hold for all vector combinations the outcome is that the three populations must conform a triangle in *S*_*f*_ with all three angles acute. We will refer to this as the *triangle criterion*. Thus, in terms of *f*_3_ the requirements for *f*_2_ to act as a true distance in *S*_*f*_ are

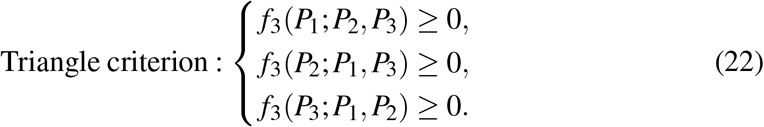

Only in that case *f*_2_ acts as a mathematical distance, the distance matrix 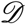 makes sense, and the results of Section 3 concerning treeness apply. A similar qualitative description appears in [4, Fig. 1.17.1].

Had we four populations to be compared, then the image would be that of a tetrahedron with all angles acute and therefore twelve inequalities in (22) that would secure *f*_2_ acting as true metric within this set of four populations.

In this context, finding one value *f* _3_ < 0 means that the assumed metric *f*_2_ has lost its distance character and thus the mathematical model that studies a phylogeny does not apply to that combination of vectors. One reason may be that these populations are rather related by an admixture process. This is illustrated in Figure 1b, where the vertex *P*_x_ in the triangle is *too close* to the segment 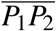; and the meaning of too close is that the angle *ϕ* at vertex *P*_x_ is obtuse. A related discussion on the distance character of *f*_2_ and formal results on additivity may be found in [12].

## 6 Applications to a treeness scenario

We review two tests used commonly in treeness cases and point out that the outcome may be conditioned by the chosen outgroup populations.

### 6.1 Outgroup *f*_*3*_–statistics test

The so–called outgroup *f*_3_–statistics test [9, 8] has been proposed to determine the closest population *P*_*i*_ (in the sense of treeness), among a set {*P*_1_, …, *P*_*n*_}, to a test population *P*_*test*_ using as reference an outgroup population *P*_*out*_. Given the tree ((*P*_*test*_, *P*_*i*_), *P*_*out*_) The longer the branch from the outgroup population *P*_*out*_ to the node (*P*_*test*_, *P*_*i*_), i.e. *f*_3_(*P*_*out*_ ; *P*_*i*_, *P*_*test*_), the closer *P*_*i*_ to *P*_*test*_. Note that the answer is not provided by the minimum value among the two–populations distances in the set {*f*_2_(*P*_*test*_, *P*_*i*_)}, with *i* = 1, …, *n*. The triangle criterion (22) should routinely precede the outgroup *f*_3_–statistics procedure in order to rule out population triplets not related by treeness.

Figure 4a illustrates this commonly used procedure. Population *P*_*a*_ is closer to *P*_*test*_ than population *P*_*b*_, because *f*_3_(*P*_*out*_ ; *P*_*test*_, *P*_*a*_) > *f*_3_(*P*_*out*_ ; *P*_*test*_, *P*_*b*_).

**Figure 4:**
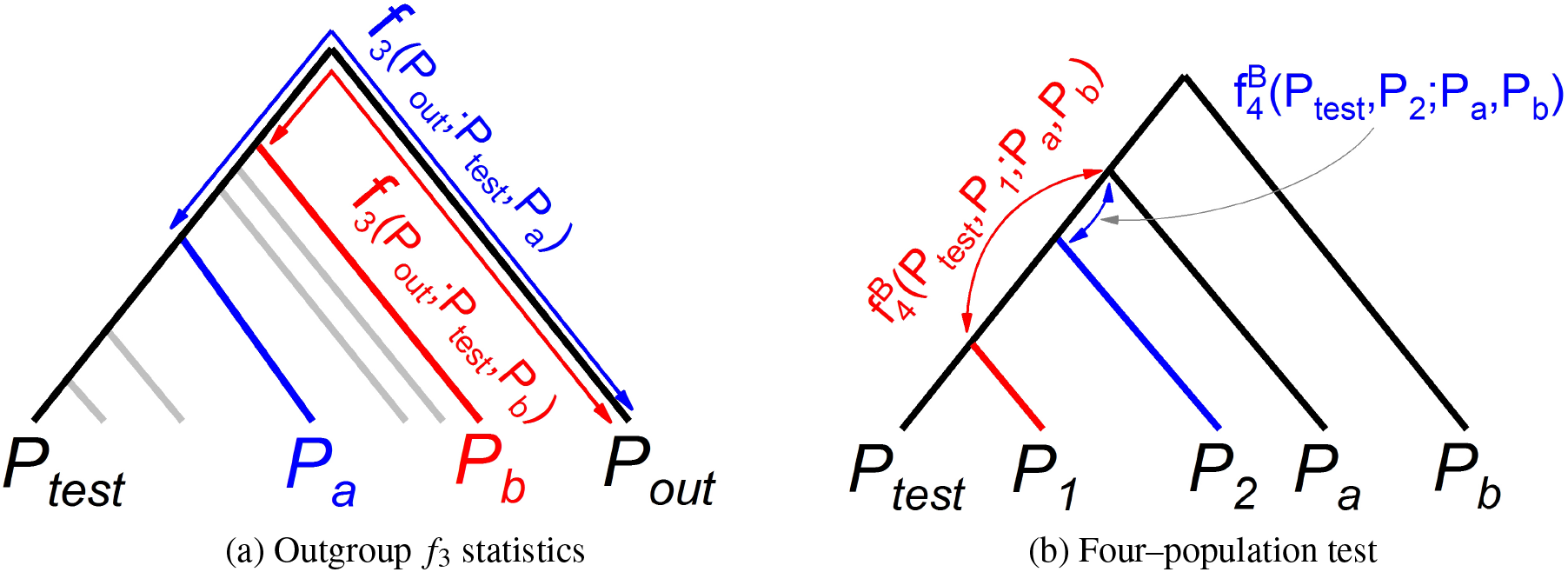
(a) Scheme for the interpretation of outgroup *f*_3_–statistics. The graph is the union of three–population trees having in common *P*_*test*_ and *P*_*out*_. *P_a_* and *P_b_* are two populations among a larger set represented by unlabelled (grey) branches. *f*_3_ gives the length of the branch between *P*_*out*_ and the nodes (*P*_*test*_, *P*_*a*_) (blue arrow) and (*P*_*test*_, *P*_*b*_) (red arrow). Here *f*_3_(*P*_*out*_ ; *P*_*test*_, *P*_*a*_) > *f*_3_(*P*_*out*_ ; *P*_*test*_, *P*_*b*_), and so *P_a_* is closer to *P*_*test*_ than *P*_*b*_. (b) Scheme for the interpretation of the four–population test. The graph is the union of two four–population trees. Coded in colour, 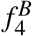 gives the inner branch length of the trees.

The outgroup *f*_3_ statistics has a drawback because the distance ranking of populations may depend on the choice of outgroup. To illustrate this we build up the following counterexample. A number of assumptions and simplifications are introduced in order to ease the explanation. It is not meant that the drawback takes place only in this singular study case.

Consider a test population *P*_*test*_, two outgroup populations 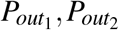, and two populations *P*_*a*_, *P*_*b*_. The question is: Which one of the latter two populations is closer to the test population? Let us assume that in space *S*_*f*_, the tips of all five population vectors are approximately on the same plane with the distribution of vector tips on such a plane as in Figure 5. This is an unlikely situation but simplifies dramatically the explanation. The only vector tip locations related via treeness with *P*_*test*_, and concomitantly with 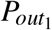 and 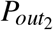, are outside the semi-circles. Tips inside the semi–circles break the triangle criterion at least with one outgroup population.

**Figure 5:**
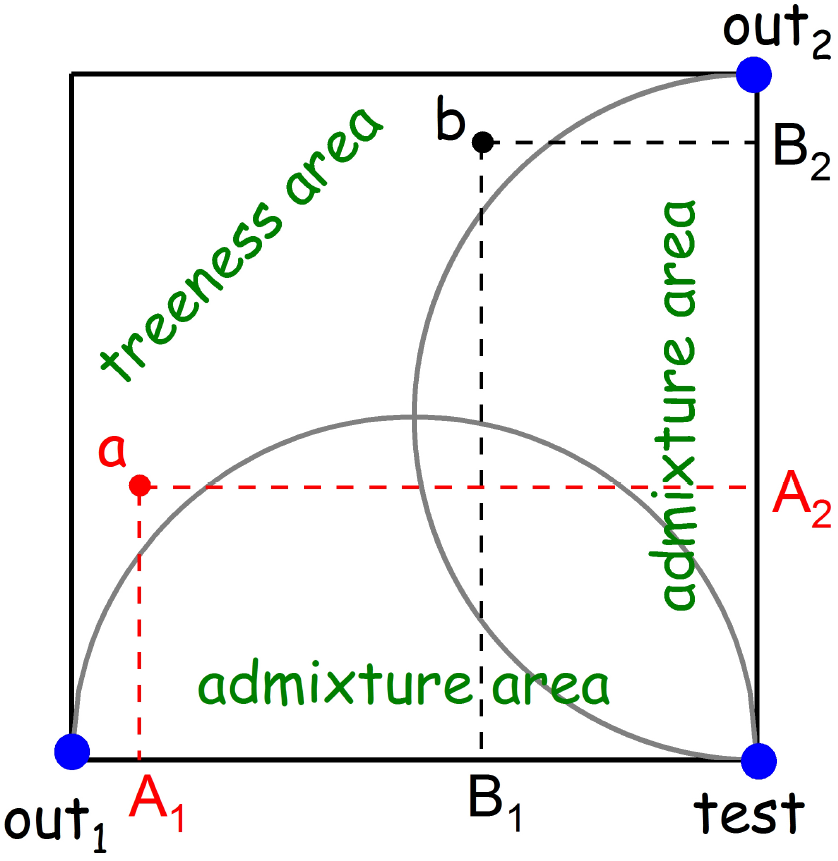
Outgroup *f*_3_–statistics test counterexample. Setup of the five population vector tips involved in. Population labels shortened for clarity.

For the sake of computation simplicity of the scalar product involved in *f*_3_, let the distances be 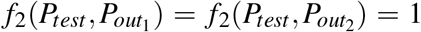, in arbitrary units. Then, using the geometric interpretation of the scalar product, the orthogonal projection of the point *P*_*a*_ onto the segment 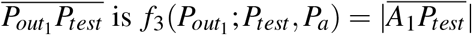. We denote the length of a segment as | · |. The three remaining projections are 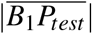, 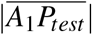 and 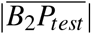.

These four quantities are related to *f*_3_ as follows:

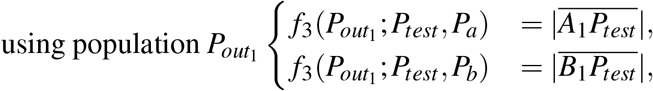

and from the graph we get 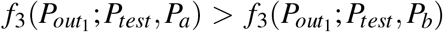. We conclude that population *P_a_* is closer to *P*_*test*_ than *P_b_*. Also

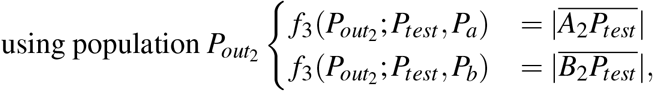

and from the graph we get 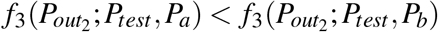. Therefore population *P*_*b*_ is closer to *P*_*test*_ than *P*_*a*_, in contradiction with the conclusion above.

This counterexample illustrates how the choice of population outgroups may affect the outcome of the test. Conclusions based on outgroup *f*_3_–statistics test should be buttressed by further arguments.

### 6.2 Four–population test

Let us replace population *P*_3_ in the outgroup *f*_3_–statistics test described above with a node so that the associated tree is (((*P*_*test*_, *P*_*i*_), *P*_*a*_), *P*_*b*_). The inner branch length is given by 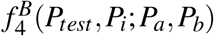. Thus, provided 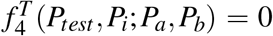, the greater the value of 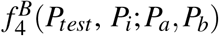 the closer the populations *P*_*i*_ and *P*_*test*_. For instance, with populations *P*_1_, *P*_2_, whenever 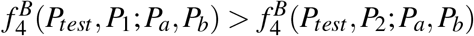, then *P*_1_ is closer to *P*_*test*_ than *P*_2_; provided: 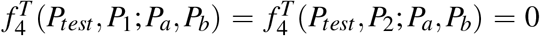, This is the instance illustrated in Figure 4b. Remind that 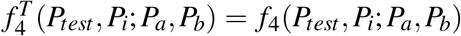, whereas 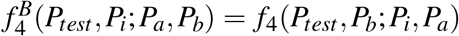.

Due to the fact that *f*_4_ can be written down as a difference of two *f*_3_ terms (see property 10 in Appendix B) the same considerations about the outgroup *f*_3_–statistics test drawback apply.

## 7 Lack of treeness: Population admixture

The lack–of–treeness case is interpreted as a population admixture scenario in *f*–statistics. The statistic *f*_2_ may be numerically computed, of course, but it has lost the interpretation as a genetic distance. Next, we first establish the algebraic description of a hybrid population, then give the population admixture criterion in terms of *f*–statistics and finally provide four different expressions to estimate the proportions of the admixture. Three of them are new results and one of them is related to the fixation index *F*_*ST*_.

### 7.1 Population admixture: Mathematical description

Let 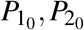 denote the parental and 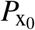 the hybrid populations at the time when the admixture took place. Let *P*_1_, *P*_2_, *P*_x_ be the evolved populations whose allele frequencies we have experimental access to. These six vectors are represented in Figure 1a in *S*_*f*_ of dimension three. In the admixture process the populations 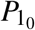 and 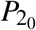 get combined to successfully form the new population 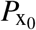. More specifically, given a particular SNP of the hybrid, its allele frequency is formed by a fraction *α* of contributor *P*_1_ and 1 − *α* of contributor *P*_2_. And similarly for all the SNPs in the array, with the same fraction. We can then write

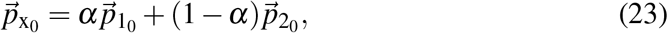

at the time the hybrid population is born. This is just the kind of linear combination vector introduced in (2). The situation illustrated in Figure 1a shows that when the ad-mixed population 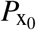 is born, the three population vectors are coplanar. Subsequently, they wander in time as a consequence of genetic evolution describing three independent crinkly trajectories in *S*_*f*_ (curly green lines). The initial segment 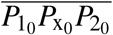 becomes triangle *P*_1_*P*_x_*P*_2_ after some time has elapsed. The goal is to estimate the fraction *α* from the three population vectors, 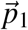, 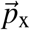, 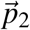. We will assume that the vector 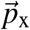 of allele frequencies of the population *P*_x_ is well approximated by the linear combination

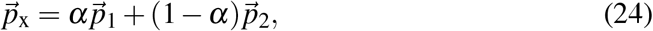

of the two contributor allele frequency vectors.

The idea of using a weighted average for one particular gene frequency of the mixed population appears already in [5, eq. (47)], [3, eq. (8.3)] and [4, eq. (1.17.1)].

Equation (24) will be a good approximation if the angle *ϕ* at vertex *P*_x_ in Figure 1b is close to 180°. *ϕ* is the separation angle between the two vectors 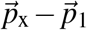 and 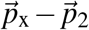, and can be computed with the aid of (3) irrespective of the dimension of *S*_*f*_

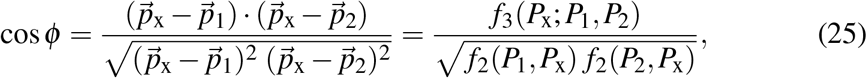

for it is given in terms of scalar products. The limiting value cos *ϕ* = − 1 corresponds to the birth of admixture. In section 5 we have discussed that cos *ϕ* < 0 corresponds to a population admixture scenario, whereas cos *ϕ* > 0 corresponds to a treeness one. The value cos *ϕ* = 0 is the separatrix of both scenarios. When cos *ϕ* = 1, the three vector tips are indeed co–linear but it is not an admixture situation because the order of vectors is not correct.

Given a set of putative contributors we might want to determine the most likely pair that generates a hybrid population. Since the function cos *ϕ* is bounded, −1 ≤ cos *ϕ* ≤ 1, it can be used to rank the contributor pairs in the set.

### 7.2 Admixture test

In *f*–statistics, whenever three populations, say *P*_1_, *P*_2_, *P*_x_, do not pass the triangle criterion for treeness the triplet is considered as candidate for population admixture. The formalism is binary in this respect. One (and only one) of the three possible *f*_3_ values is negative; say, *f*_3_(*P*_x_; *P*_1_, *P*_2_) < 0. This is the so–called *f*_3_ admixture test. We prove that *P*_x_ may be considered as a population admixture of *P*_1_ and *P*_2_.

Let us compute *f*_3_(*P*_x_; *P*_1_, *P*_2_) with *P*_x_ the hybrid described by the allele frequency vector 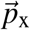 in the linear combination (24)

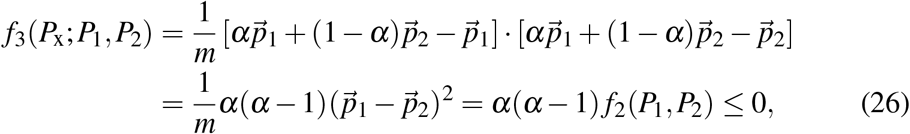

because *f*_2_ ≥ 0, and 0 ≤ *α* ≤ 1 by hypothesis of population admixture. A similar computation yields

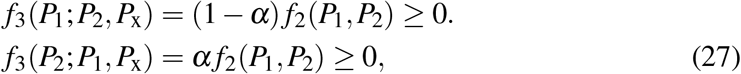

and so the two remaining *f*_3_ values are positive.

We can summarize the conditions for population admixture without explicit mention to *α*. For a population of the form (24)

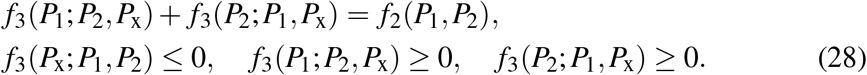

For fixed populations *P*_1_ and *P*_2_, the term *f*_2_(*P*_1_, *P*_2_) in (26) is a constant and the functional dependency between *f*_3_ and the admixture proportion *α* is given by the parabola in Figure 6 (solid black curve). Interestingly, in [8, Figure 5B] a similar parabolic curve is obtained from a numerical simulation in which *f*_3_ has been expressed in terms of expected coalescence times and parameters have been chosen so that *f*_3_ < 0. A few measurements by hand on that plot yielded the dashed curve with the shaded area as confidence interval in Figure 6.

**Figure 6:**
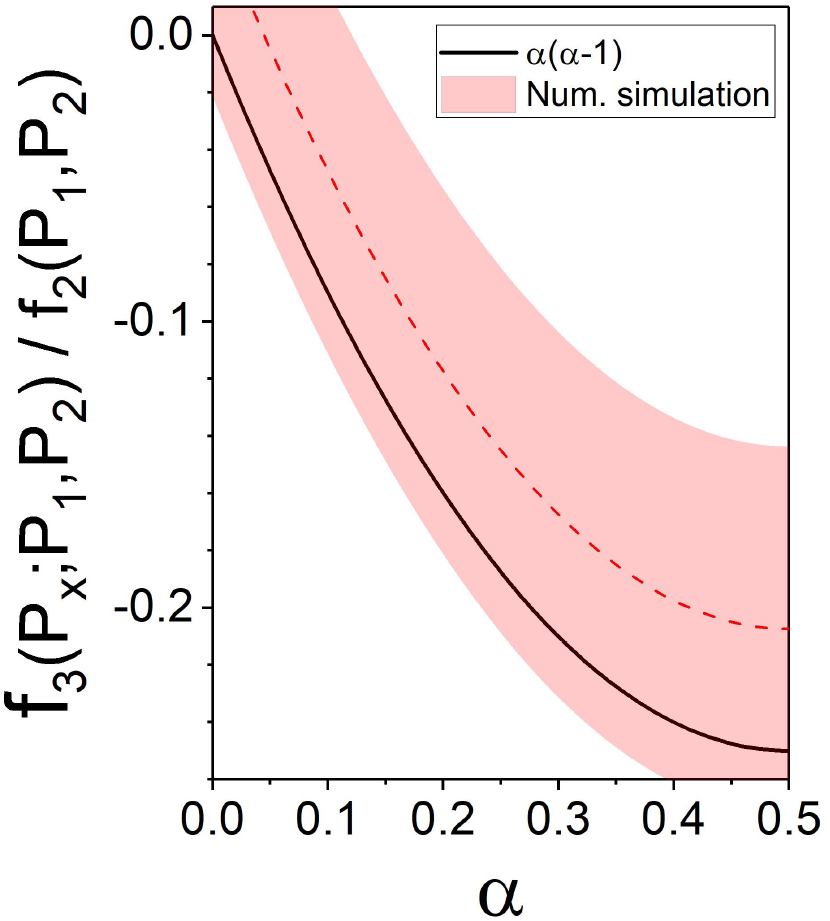
Relationship between *f*_2_, *f*_3_, and the admixture proportion *α* according to (26) (black curve). Numerical simulation taken from [8, Fig.5B] (dashed red) with confidence interval (shaded).

### 7.3 Population admixture: Determination of proportions

Given the population vectors 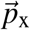, 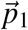, 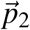, four formulas are given to estimate the proportion *α* in (24). The first one, based on a least squares procedure, is new. The second one relies on orthogonal projections on an arbitrary direction in *S*_*f*_ determined by two auxiliary populations and gives rise to the so–called *f*_4_–ratio statistics. The last two ones are simplifications using only one auxiliary population and are new.

#### 7.3.1 Approach 1: Least squares

The least squares solution to (23) is readily obtained using vector algebra. In this simple case no matrix algebra is needed. Let us rewrite (24) as

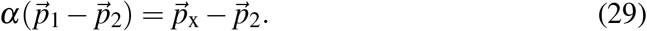

The scalar product by 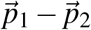 of both sides yields

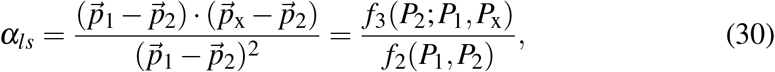

which is the least squares estimate for *α*. The conventional confidence interval reads

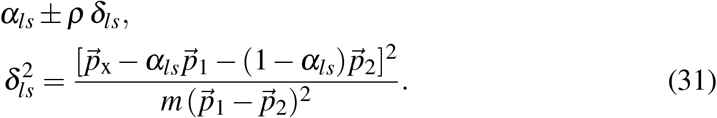

For a 95% confidence interval and *m* large enough, *ρ* = 1.96.

Figure 1b provides the geometric interpretation of the leftmost fraction in (30). The denominator is a squared vector modulus: 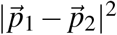. According to the geometric interpretation of the scalar product 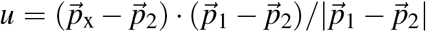 is the length of the projection of segment 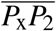 along the direction of vector 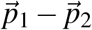, and therefore 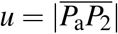. The estimate in (30) is then the ratio *α_ls_* = *u/v*, with 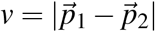. At the time population *P*_x_ emerges, i.e. 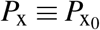, the estimate is exact.

#### 7.3.2 Approach 2: Projection in *S*_*f*_ using two auxiliary populations

Consider two arbitrary populations *P*_*a*_ and *P*_*b*_. The difference vector 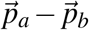 defines an arbitrary direction in *S*_*f*_. The scalar product by 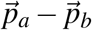 of both sides in (29) gives an estimate involving five populations

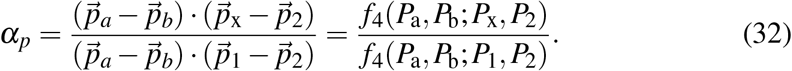

According to the geometric interpretation of the scalar product given in Section 2, the numerator is a length proportional to the orthogonal projection of vector 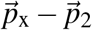 along the direction defined by vector 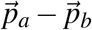. Similarly for the denominator with the difference vector 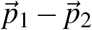. The estimate *α*_*p*_ is then the ratio of both lengths, provided the denominator is non–vanishing. This result is referred to as *f*_4_–ratio statistic and populations *P*_*a*_, *P*_*b*_ are outgroups. Notice that, from an algebraic point of view, there is no constraint on the choice of populations *P*_*a*_, *P*_*b*_. The sub–index in *α_p_* refers to the orthogonal projection involved in the method. Since we are not in a treeness scenario, no argument permutation is made in the computation of *f*_4_.

The interpretation of (32) is in Figure 7 where the five populations are represented by their allele frequency vectors in dimension three. The vector 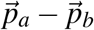 defines an arbitrary direction in *S*_*f*_. Segments 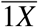 and 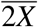 are projected onto 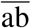. The length ratio 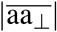 over 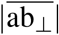 is *α*_*p*_ in (32). In practice, *α*_*p*_ is computed for a number of population pairs *P*_*a*_, *P*_*b*_, which therefore define different directions in *S*_*f*_. After that, a re–sampling procedure provides a confidence interval [7].

**Figure 7:**
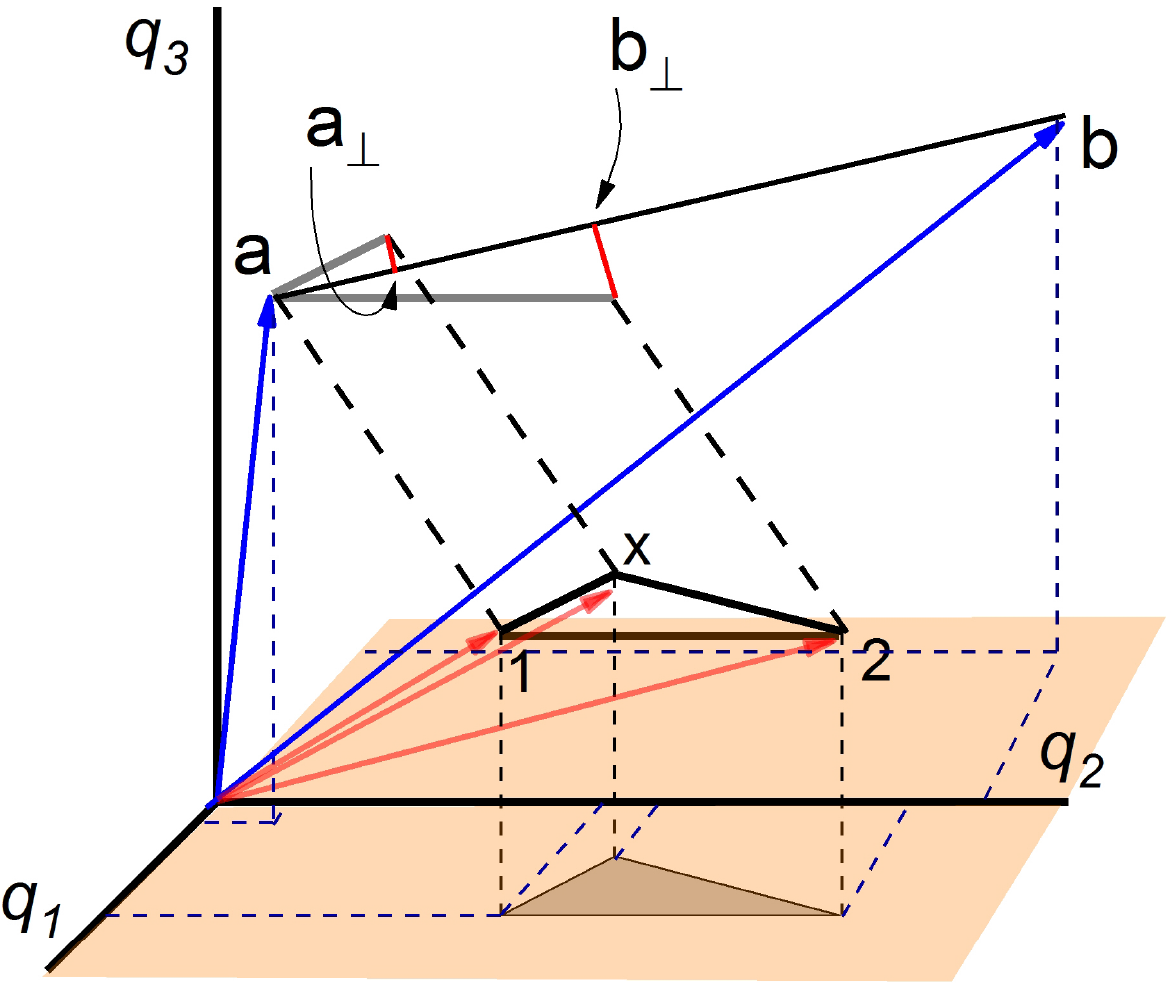
Geometric representation of the elements involved in the admixture *f*_4_–ratio statistics (32) for *S*_*f*_ of dimension three. *P*_a_ and *P*_b_ are reference populations. *P*_x_ is studied as a possible mixture of *P*_1_ and *P*_2_. The five frequency allele vectors have been drawn (red). In the triangle conformed by populations {*P*_1_, *P*_2_, *P*_x_} the angle at vertex *P*_x_ is obtuse. The projections on the horizontal plane are just for helping with the perspective. Segments 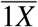 and 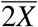 have been parallel transported to point a in order to illustrate their orthogonal projection onto the line 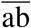. The quotient of length 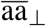 over length 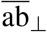 is just *α*_*p*_ in (32). Population labels shortened for clarity.

#### 7.3.3 Approach 3: Projection in *S*_*f*_ using one auxiliary population

If in (32) we choose either *P*_*b*_ = *P*_1_ or *P*_*b*_ = *P*_2_ we get two further formulas involving only one arbitrary auxiliary population

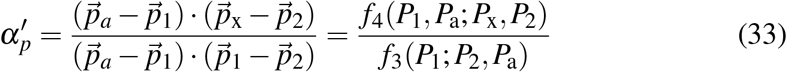

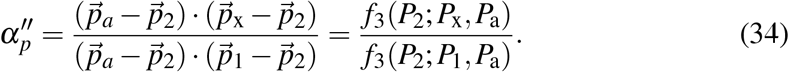

These equations may be seen as a simplification the *f*_4_–ratio (32) in the sense that only one auxiliary population is involved.

### 7.4 An illustrative example

In order to grasp the similarities and differences among the estimates *α*_*ls*_, *α*_*p*_, 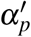 and 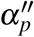, we have carried out the following numerical simulations.

From the population dataset *v37.2.1240K HumanOrigins* from *Reich Lab* [13] we have taken two parental populations, *P*_1_ =*English* and *P*_2_ =*Yoruba*, and build up the admixed population (23), 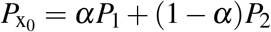 for several values of *α*.

To simulate finite time evolution we have added some noise of amplitude *ε* to the 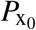 allele frequencies. The rationale is that the greater the *ε* value the longer the time elapsed since population admixture event. The parent populations stay invariant in the simulation for the sake of simplicity. The idea is just to take the vector 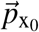 off the plane defined by the two contributor vectors. Besides, nine auxiliary populations have been chosen from the same dataset, namely: Han, Papuan, Surui, Karitiana, Aleut, Ju hoan North, Gujarati, Iranian and Australian. The simulations have been carried out as follows.

The time evolved hybrid *P*_x_ is simulated for *α* = 0.1, 0.2,…, 0.5, and *ε* = 0.1,…, 0.4. The outcomes of the simulations are in Figures 8 and 9.

**Figure 8:**
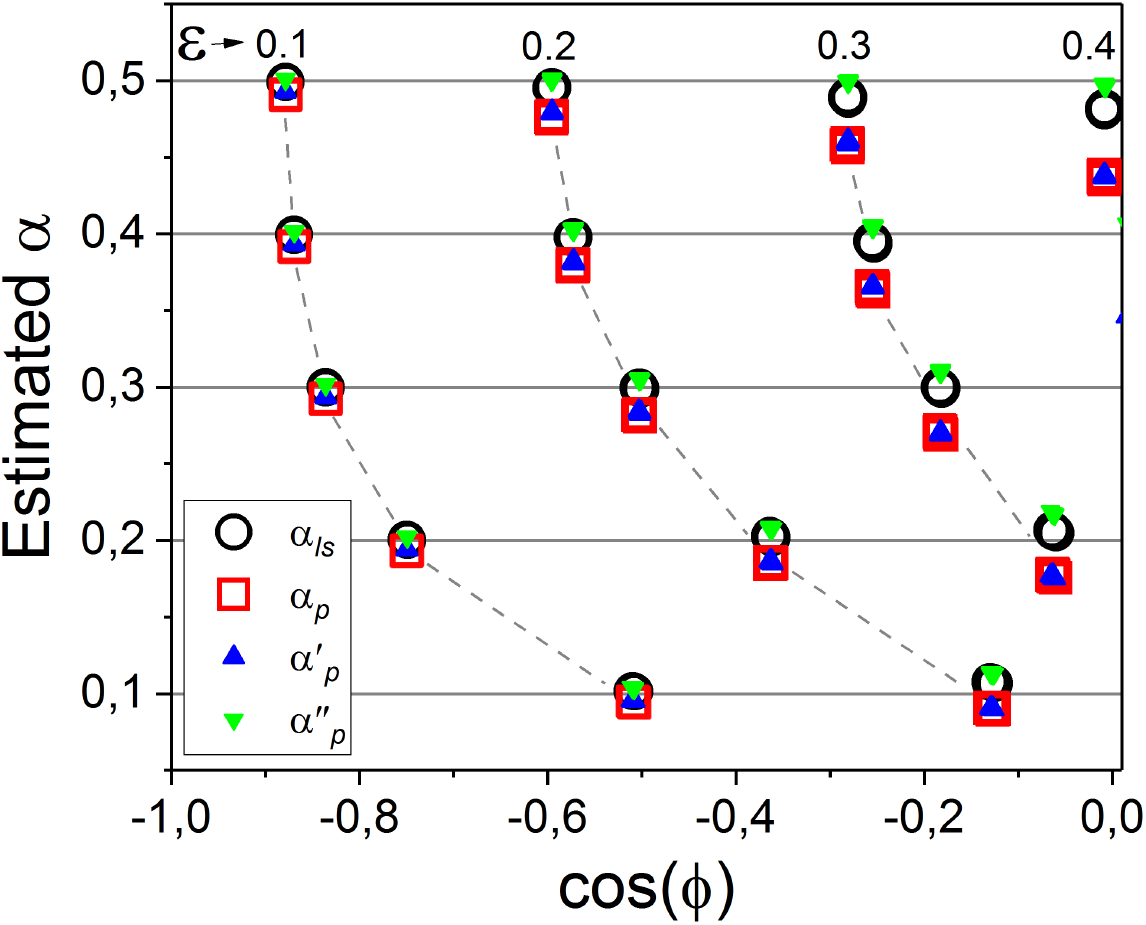
Estimates of *α*_*ls*_ in (30), *α*_*p*_ in (32), 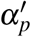 in (33) and 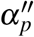 in (34), as a function of cos *ϕ* in (25) for five different values of *α* used to generate the hybrids, which are indicated by the horizontal grid lines. The *ε* value is related to the time since the hybridization event: the greater the *ε* value the more time elapsed since the mixing event. Dashed lines are only for visual help.

**Figure 9:**
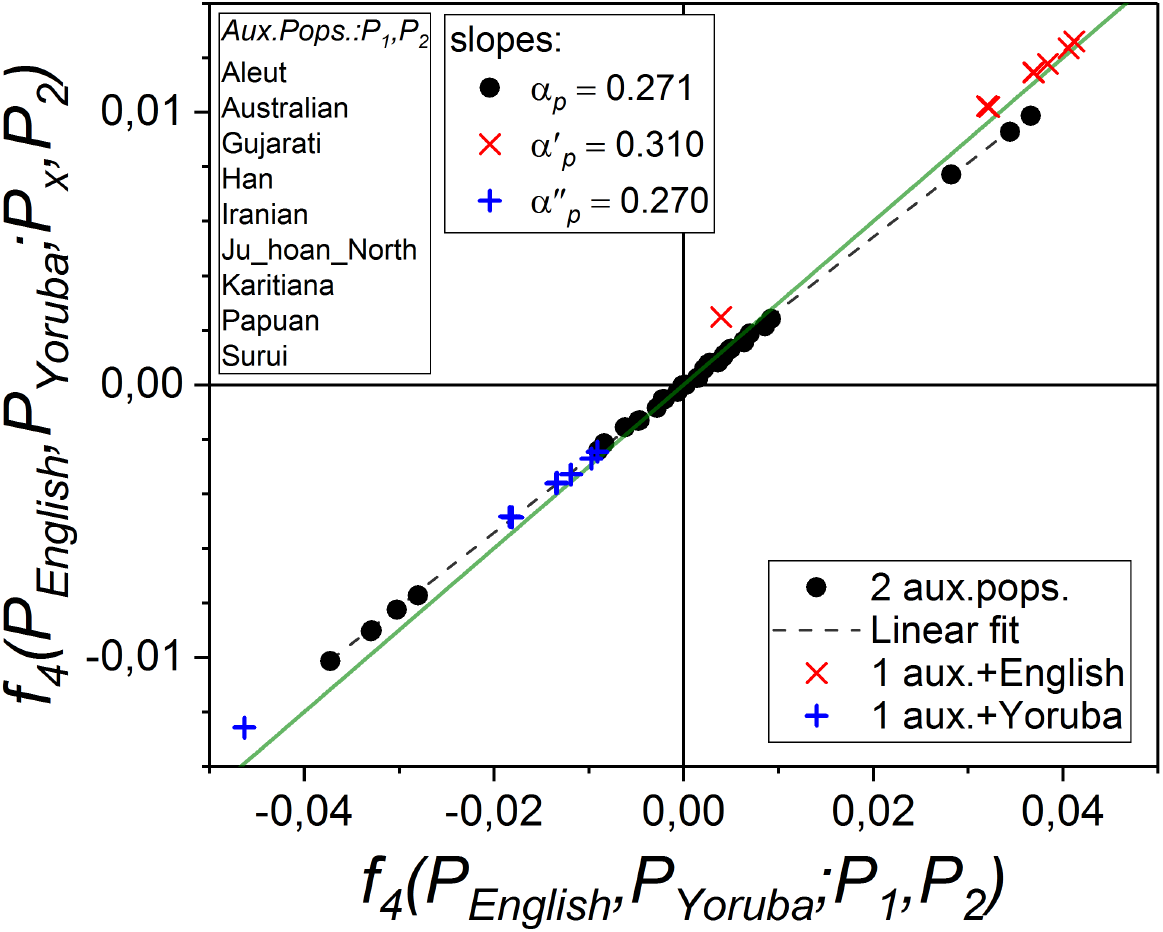
*f*_4_–ratio plane for a simulation with initial *α* = 0.3 in (23). The axes of the plot represent the numerator and denominator of (32). Black dots stand for different evaluations with the 36 outgroup pairs from the boxed list. The slope of the linear fit (dashed line), with fix intercept at the origin, is the average estimate for *α_p_* in (32). Fit lines to pluses and crosses (not showed) give averages for 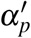 in (33) and 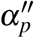 in (34), which involve only one auxiliary population. The solid green line has slope *α* = 0.3 and is given for the sake of comparison.

As regards *Approach 1*, given the three populations *P*_1_, *P*_2_, *P*_x_, equation (30) determines univocally *α*_*ls*_. In Figure 8 the open circles stand for the *α*_*ls*_ estimates as a function of the cos *ϕ* value reached. Irrespective of the *ε* value, the error estimate (31) is in the range 0.004 < *ρδ*_*ls*_ < 0.008. Remind that −1 < cos *ϕ* < 0 is the admixture condition and cos *ϕ* = −1 at admixture birth. For fixed *ε* value, an unbalanced admixture heads toward a genetic drift scenario (cos *ϕ* > 0) faster than a balanced one in this type of simulation.

To deal with *Approach 2* we read (32) as a linear relationship between 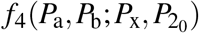) and 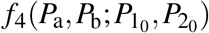), instead of a ratio. Then a plot of pairs of both quantities, evaluated for different populations *P*_*a*_, *P*_*b*_, should appear as points on a straight line. This behaviour is observed in Figure 9 (black dots) as well as for the other *α* values used in the simulation. In the present case the slope of the fit (dashed line), which was constrained to intercept the origin, gives *α*_*p*_ = 0.271 ± 0.001. The solid line has slope 0.3 and is given for the sake of comparison. The whole set of *α*_*p*_ estimates is in Figure 8 (open squares).

The outcomes of *Approach 3*, which involves only one auxiliary population, are also in Figure 9. Red pluses and crosses stand for the cases where *P*_*English*_ or *P*_*Yoruba*_ act as auxiliary population, respectively. The estimated slopes are 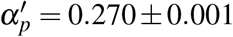 and 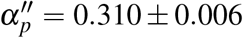 (fit line not showed). Further results are in Figure 8 (solid triangles).

## 8 Population admixture: Coalescence times, pairwise sequence differences and fixation index *F*_*ST*_

### 8.1 Coalescence times

A result reported in [8]. provides an alternative admixture proportion estimate on the basis that *f*_2_(*P*_*i*_, *P*_*j*_) can be written in terms of expected coalescence times *T*_*ij*_ of populations *P*_*i*_ and *P*_*j*_ to a common ancestor *P*_*a*_. The formula in [8] is given under the assumption that all populations involved are sampled at the same time

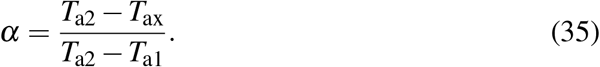

According to [8], when the number of individuals in populations is large

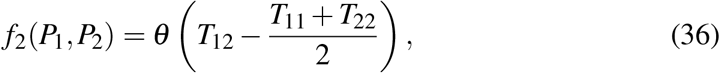

with *θ* / 2 the constant rate of mutation on each branch of the tree. Writing explicitly down the *f*_4_–ratio (32) in terms of *f*_2_, and according to (13) (see also B)

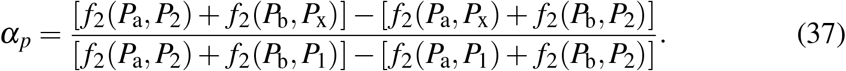

Substituting (36)

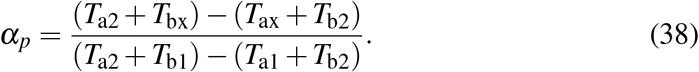

Both formulas are not so different if the sums in parentheses are interpreted as averages of two coalescence times.

### 8.2 Pairwise sequence differences

A second outcome in [8] is based on the relationship between *f*_2_ and the average number of pairwise sequence differences *π*_*ij*_ between populations *P*_*i*_ and *P_j_* which reads

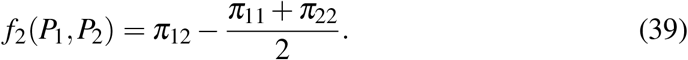

Substitution in (37) yields

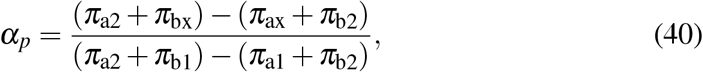

to be compared with the formula in [8]

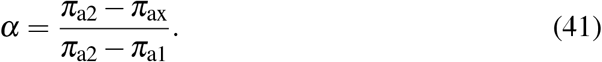

The comparison is similar to that of coalescence times. According to numerical simulations in [8], both estimates seem to perform well although for low admixture proportion, the latter performs better.

### 8.3 Fixation index *F*_*ST*_

In [4, Sec.1.17], the populations are referred to a multidimensional *gene–frequency space* that clearly plays a role similar to *S*_*f*_. Moreover, the triangle criterion is qualitatively described and the authors provide, without disclosing, a formula for the population admixture proportion in terms of the so–called *F*_*ST*_ genetic distance. We prove that such a formula and *α*_*ls*_ in (30) are related results.

The formula in [4] for the admixture proportion *m* reads

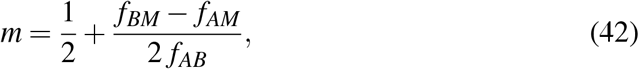

where *f*_*AB*_ stands for the *F*_*ST*_ distance between contributors *A* and *B*; and *f*_*AM*_, *f*_*BM*_, the distances between each of the two contributors and the hybrid *M*. Here *m* is the admixture proportion of population *A*. In the case of two populations, the difference between *F*_*ST*_ and *f*_2_ is only an approximately constant normalization factor. Consequently, the formula (42) can be re–written as

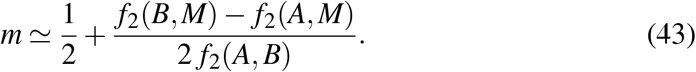

Now, using (16) we get

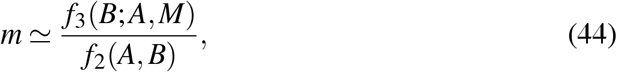

and, by (30), the conclusion *m* ≃ *α*_*ls*_ follows.

## 9 Discussion and conclusions

We have introduced the *f*–statistics as the set of constraints needed for a genetic distance matrix to be associated to a phylogenetic tree of four populations. As far as *f*_2_ can be interpreted as a true genetic distance in *S*_*f*_, the concomitant scenario is tree–like–ness. Else, the case is interpreted as population admixture.

Populations are described as vectors in *S*_*f*_ evolving in time. A long enough time lapse may drive an admixture scenario into an evolutionary one, and the way around. It suffices that, as a consequence of evolution, the wandering of population vectors transforms an obtuse–angled triangle in *S*_*f*_ into an acute–angled one, or viceversa. An instance of this class [7, 8] points out that high population–specific drift in the admixed population could prevent a negative value for the *f*_3_ test. In Appendix A we obtain a conservative formal lower bound to the linear distance that vectors in *S*_*f*_ may travel before a population admixture can be detected as an evolutionary process.

The outgroup *f*_3_–statistics test and the four–population test rank a population set according to a genetic distance. These tests should be preceded by the triangle criterion in order to rule out populations not related by an evolutionary process. We have noted that contradictory rankings may appear between outcomes obtained with different outgroups.

The four formulas provided to estimate the proportion of contributors in a hybrid population rely on the use of a linear combination of population vectors in *S*_*f*_. This idea can be traced back to [4] where the triangle criterion is already qualitatively described. The new estimate (30) is obtained from an orthogonal projection on a fixed direction defined by the admixture contributors. It can be also interpreted as a least squares approximation and is related to a result based on the *F*_*ST*_ genetic distance. The way we have derived the formula for the *f*_4_–ratio (32) is based on orthogonal projections onto an arbitrary direction in *S*_*f*_ (defined by the difference vector of the two auxiliary populations). The two remaining new formulas are obtained by similar projections in *S*_*f*_ but using only one auxiliary population. The simulations carried out illustrate the behaviour of the four estimates. It seems that *α*_*ls*_ gives balanced estimates whereas *α*_*p*_ and 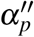 tend to underestimate and 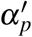 to overestimate, albeit more systematic simulations are needed to base these conclusions.

The *f*–statistics formalism has introduced an important tool to facilitate the study of intricate population structures. We have presented a new derivation from a geometry–based scheme, as well as some new results and relationships with other formalisms, that may ease the use and interpretation of the *f*–formalism in practice.

## Acknowledgements

We are indebted to J.M. Bordes for illuminating discussions about the geometry of *f*–statistics. GOG thanks The Leverhulme Trust for financial support. JAO work was partially supported by the Spanish MINECO (grant numbers AYA2016-81065-C2-2 and PID2019-109592GB-100).

## A Lower bound to reliability of *f*_3_ admixture test

A lower bound may be given to the Euclidean distance *r* travelled in *S*_*f*_ by three points representing an admixed population and two contributors, respectively, before they reach the right–angled triangle geometric configuration. If the rate at which allele frequencies change in the dataset were known, then an estimate of the time *t* elapsed since the birth of the admixture could be provided on the basis that *r* is proportional to 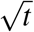, if we assume that the vectors describe Brownian trajectories in *S*_*f*_. These bounds are of interest to asses the validity of the *f*_3_ admixture test.

In Figure 10, the point *A′* represents the admixture population and *A* and *A″* stand for the two equal–weight contributors to the linear combination that defines de birth of *A′*. The goal is to give just a lower bound, so we do not deal with a generic proportion. Besides, *A* and *A″* are assumed one unit apart and so define the genetic distance scale. In *S*_*f*_ those points evolve in time describing a Brownian–like trajectory (green lines) and the linear distance reached *r*, is proportional to 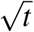. After a lapse of time, the trajectory born at initial time at *A* wanders in *S*_*f*_ until it crosses the circumference at *B*, and at a later time it will reach the point *C*.

**Figure 10:**
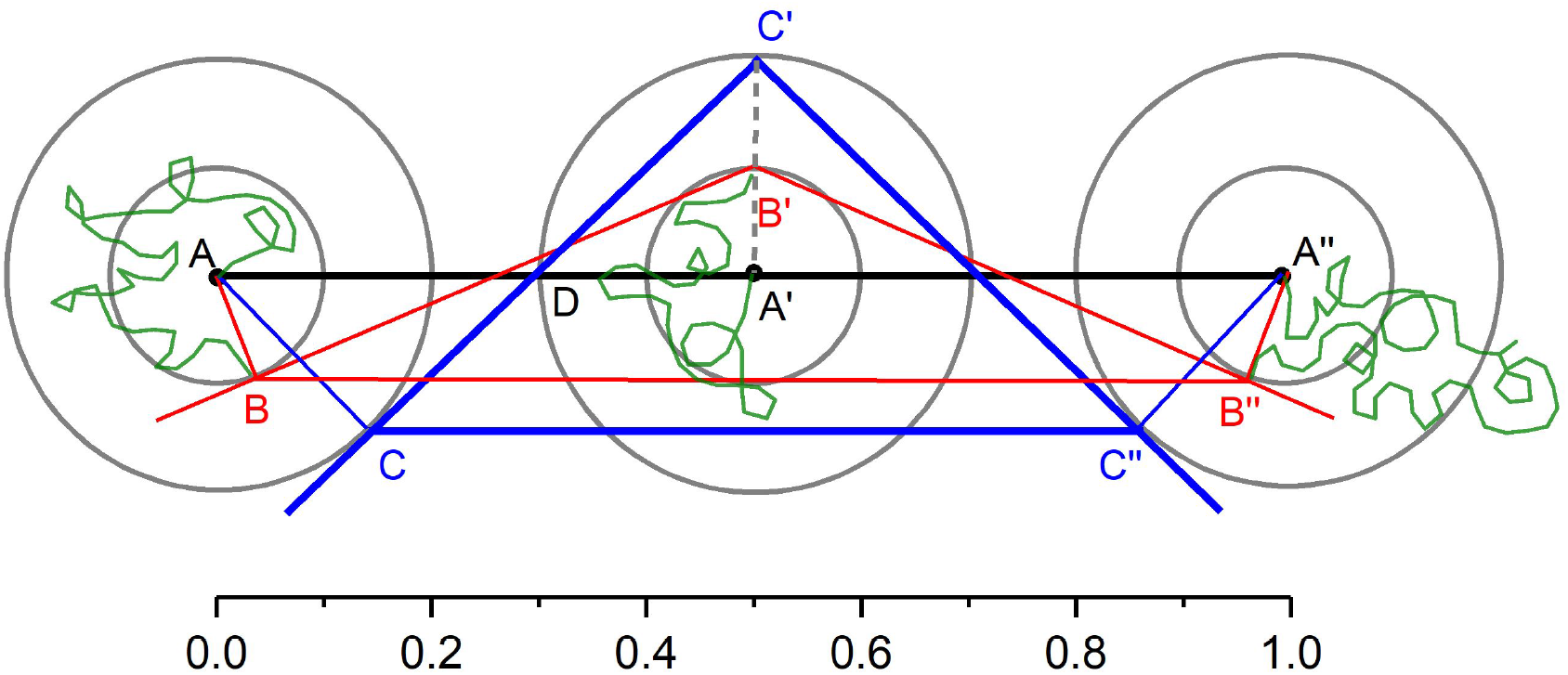
Scheme to find a lower bound to reliability o f *f*_3_ admixture test.

We are interested in the situation where, given a radius, the angle at vertex *B′* is as small as possible. We assume the favourable case where the positions *B*, *B′*, *B″*, are simultaneously reached. This is the configuration depicted, where lines 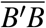 and 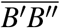,, are tangents to a circumference. This is only an intermediate situation. We want to ascertain the radius *r* at which the angle at vertex *B′* becomes right as in *C′*. The proof relies on the fact that the triangles *ACD* and *DA′C′* are identical. Thus, 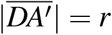 and 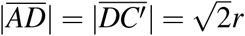. Since 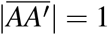 we have then 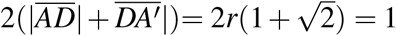, and therefore 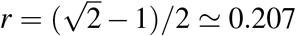. Of course, this picture corresponds to an extreme situation in which the wandering of the three trajectories ends up simultaneously at points *C*, *C′* and *C″*. This is why 21% of the initial distance between contributors is just a lower bound. After that, if the dynamics leads the geometry to an acute-angled triangle the *f*_3_ admixture test will fail.

An algebraic proof is also possible minimizing the expression of *r* with respect to several variables but the algebra is quite more involved. It yields the very same result for the radius *r*.

## B Some useful algebraic identities

A listing of algebraic identities among statistics may be useful. The following one does not exhaust the list of possible relationships. All can be straightforwardly proved, for instance by expressing *f*_3_ and *f*_4_ in terms of *f*_2_. Some of them are mere symmetry properties under exchange of arguments.

1. *f*_2_(*P*_1_, *P*_2_) = *f*_2_(*P*_2_, *P*_1_).
2. *f*_3_(*P*_*a*_; *P*_1_, *P*_2_) = *f*_3_(*P*_*a*_; *P*_2_, *P*_1_).
3. 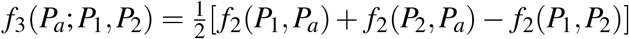.
4. *f*_3_(*P*_1_; *P*_2_, *P*_*a*_) + *f*_3_(*P*_2_; *P*_1_, *P*_*a*_) = *f*_3_(*P*_1_; *P*_2_, *P*_*b*_) + *f*_3_(*P*_2_; *P*_1_, *P*_*b*_).
5. *f*_4_(*P_a_, P_b_*; *P*_1_, *P*_2_) = *f*_4_(*P*_1_, *P*_2_; *P_a_, P_b_*) = *f*_4_(*P_b_, P_a_*; *P*_2_, *P*_1_).
6. *f*_4_(*P_a_, P_b_*; *P*_1_, *P*_2_) = − *f*_4_(*P_a_, P_b_*; *P*_2_, *P*_1_) = − *f*_4_(*P_b_, P_a_*; *P*_1_, *P*_2_).
7. 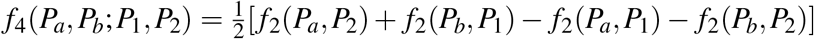.
8. *f*_4_(*P*_1_, *P*_2_; *P*_3_, *P*_4_) = *f*_4_(*P*_1_, *P*_3_; *P*_2_, *P*_4_) + *f*_4_(*P*_1_, *P*_4_; *P*_3_, *P*_2_).
9. *f*_3_(*P*_1_; *P*_2_, *P*_*b*_) + *f*_3_(*P*_2_; *P*_1_, *P*_*a*_) = *f*_2_(*P*_1_, *P*_2_) + *f*_4_(*P_a_, P_b_*; *P*_1_, *P*_2_).
10. *f*_4_(*P_a_, P_b_*; *P*_1_, *P*_2_) = *f*_3_(*P*_*a*_; *P_b_, P*_2_) − *f*_3_(*P*_*a*_; *P_b_, P*_1_) = *f*_3_(*P*_*b*_; *P_a_, P*_1_) − *f*_3_(*P*_*b*_; *P_a_, P*_2_).
11. *f*_4_(*P_a_, P_b_*; *P*_1_, *P*_2_) = *f*_4_(*P_a_, P_b_*; *P*_1_, *P*_3_) + *f*_4_(*P_a_, P_b_*; *P*_3_, *P*_2_)

They identities above are valid in both evolutionary and population admixture contexts. In an admixture scenario: 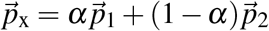, we have

1. *f*_3_(*P*_x_; *P*_1_, *P*_2_) = *α*(*α* − 1) *f*_2_(*P*_1_, *P*_2_) ≤ 0.
2. *f*_3_(*P*_1_; *P*_2_, *P*_x_) = (1 − *α*) *f*_2_(*P*_1_, *P*_2_) ≥ 0.
3. *f*_3_(*P*_2_; *P*_1_, *P*_x_) = *α f*_2_(*P*_1_, *P*_2_) ≥ 0.
4. *f*_3_(*P*_1_; *P*_2_, *P*_x_) + *f*_3_(*P*_2_; *P*_1_, *P*_x_) = *f*_2_(*P*_1_, *P*_2_).
5. *α* = *f*_4_(*P*_*a*_, *P*_*b*_; *P*_x_, *P*_2_)/ *f*_4_(*P*_*a*_, *P*_*b*_; *P*_1_, *P*_2_).
6. *f*_4_(*P*_*a*_, *P*_*b*_; *P*_*x*_, *P*_2_) = *α f*_4_(*P*_*a*_, *P*_*b*_; *P*_1_, *P*_2_).
7. *f*_4_(*P*_*a*_, *P*_*b*_; *P*_*x*_, *P*_*c*_) = *α f*_4_(*P*_*a*_, *P*_*b*_; *P*_1_, *P*_*c*_) + (1 − *α*) *f*_4_(*P*_*a*_, *P*_*b*_; *P*_2_, *P_c_*).

Eventually, remind that *f*_4_ has a twofold role in a phylogenetic scenario:

1. 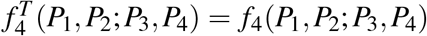
2. 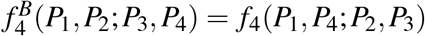

## References

[1] J.P. Barthelemy and A. Guenoche. Trees and proximity representations. Wiley, 1991.

[2] L.L. Cavalli-Sforza. Population structure and human evolution. Proceedings of the Royal Society of London. Series B, Containing papers of a Biological character. Royal Society (Great Britain), 164:362–379, 1966.

[3] L.L. Cavalli-Sforza and W.F. Bodmer. The genetics of human populations. Dover Publications, 1999.

[4] L.L. Cavalli-Sforza, P. Menozzi, and A. Piazza. The history and geography of human genes. Princeton University Press, 1994.

[5] L.L. Cavalli-Sforza and A. Piazza. Analysis of evolution: Evolutionary rates, independence and treeness. Theoretical Population Biology, 8:127–165, 1975.

[6] A.M. Harris and M. DeGiorgio. Admixture and Ancestry Inference from Ancient and Modern Samples through Measures of Population Genetic Drift. Human Biology, 89:21, 2017.

[7] N. Patterson, P. Moorjani, Y. Luo, S. Mallick, N. Rohland, Y. Zhan, T. Genschoreck, T. Webster, and D. Reich. Ancient admixture in human history. Genetics, 192:1065–93, 2012.

[8] B.M. Peter. Admixture, Population Structure, and F-Statistics. Genetics, 202:1485–501, 2016.

[9] M. Raghavan, P. Skoglund, K.E. Graf, M. Metspalu, A. Albrechtsen, I. Moltke, S. Rasmussen, T.W. Stafford Jr, L. Orlando, E. Metspalu, M. Karmin, K. Tambets, S. Rootsi, R. Mägi, P.F. Campos, E. Balanovska, O. Balanovsky, E. Khusnutdinova, S. Litvinov, L.P. Osipova, S.A. Fedorova, M.I. Voevoda, M. DeGiorgio, T. Sicheritz-Ponten, S. Brunak, S. Demeshchenko, T. Kivisild, R. Villems, R. Nielsen, M. Jakobsson, and E. Willerslev. Upper Palaeolithic Siberian genome reveals dual ancestry of Native Americans. Nature, 505:87–91, 2014.

[10] D. Reich, K. Thangaraj, N. Patterson, A.L. Price, and L. Singh. Reconstructing Indian population history. Nature, 461:489–494, 2009.

[11] J.M.S. Simões-Pereira. A note on the tree realizability of a distance matrix. Journal of Combinatorial Theory, 6:303–310, 1969.

[12] S. Soraggi and C. Wiuf. General theory for stochastic admixture graphs and F-statistics. Theoretical Population Biology, 125:56–66, 2019.

[13] David Reich Lab (Harvard University). https://reich.hms.harvard.edu/downloadablegenotypes-present-day-and-ancient-dna-data-compiled-published-papers. 2020.

[14] S. Wright. The genetical structure of populations. Annals of Eugenics, 15:323–354, 1951.

